# SARS-CoV-2 viral budding and entry can be modeled using virus-like particles

**DOI:** 10.1101/2020.09.30.320903

**Authors:** Caroline B. Plescia, Emily A. David, Dhabaleswar Patra, Ranjan Sengupta, Souad Amiar, Yuan Su, Robert V. Stahelin

**Affiliations:** Department of Medicinal Chemistry & Molecular Pharmacology, Purdue Institute of Inflammation, Immunology, and Infectious Disease, Purdue University, West Lafayette, IN 47907, United States

**Keywords:** coronavirus, electron microscopy (EM), Golgi, membrane bilayer, SARS-CoV-2, virus-like particle (VLP), viral protein, virology, virus, virus assembly, virus budding, virus entry

## Abstract

Severe acute respiratory syndrome coronavirus 2 (SARS-CoV-2) was first discovered in December 2019 in Wuhan, China and expeditiously spread across the globe causing a global pandemic. While a select agent designation has not been made for SARS-CoV-2, closely related SARS-CoV-1 and MERS coronaviruses are classified as Risk Group 3 select agents, which restricts use of the live viruses to BSL-3 facilities. Such BSL-3 classification make SARS-CoV-2 research inaccessible to the majority of functioning research laboratories in the US; this becomes problematic when the collective scientific effort needs to be focused on such in the face of a pandemic. In this work, we assessed the four structural proteins from SARS-CoV-2 for their ability to form viruslike particles (VLPs) from human cells to form a competent system for BSL-2 studies of SARS-CoV-2. Herein, we provide methods and resources of producing, purifying, fluorescently and APEX2-labeling of SARS-CoV-2 VLPs for the evaluation of mechanisms of viral budding and entry as well as assessment of drug inhibitors under BSL-2 conditions.

## Introduction

Severe acute respiratory syndrome SARS-coronavirus 2 (SARS-CoV-2) emerged in December 2019 in Wuhan, China and has since spread around the globe. As of late September 2020, the virus has been detected in 214 different countries and territories with 33 million globally confirmed cases and more than one million attributed fatalities [**1**]. The virulence of coronaviruses has previously been observed in SARS-CoV-1 and Middle East Respiratory Syndrome coronavirus (MERS-CoV) outbreaks in the previous two decades; however, there still remains no FDA-approved vaccine or treatment for *any* coronavirus. In order to develop vaccines and therapeutics, the ability to study viruses must be accessible. Under current circumstances, the authentic live SARS-CoV-2 virus is restricted to BSL-3 containment facilities; while many of these facilities exist and have refocused their collective efforts on SARS-CoV-2 research, the greater scientific community could be vastly helpful in combating this pandemic if such research was BSL-2 compatible.

For instance, BSL-2 models of other difficult-to-work-with BSL-3 and −4 pathogens such as SARS-CoV-1 [**2**], MERS [**3**], Ebola virus [**4,5**] Marburg virus [**5, 6**], and Lassa virus [**7**] have been implemented in the form of virus-like particles (VLPs). Thus, the development of BSL-2 compatible models and assays to study SAS-CoV-2 assembly, budding, and entry, as well as evaluate potential therapeutics is imperative. In this work, we aimed to develop morphologically and functionally relevant BSL-2 compatible VLPs to model SARS-CoV-2 budding and entry.

SARS-CoV-2 has a positive-sense single-stranded RNA genome of 29.7 kilobases which shares 79.6% sequence identity with SARS-CoV-1 [**8**]. Both SARS-CoV-1 and SARS-CoV-2 utilize host cell surface receptor Angiotensin-converting enzyme 2 (ACE2) to stimulate cellular uptake of bound viral particles [**8**]. Trafficked through the endocytic system of the cells, SARS-CoV-2 is eventually released into the cytoplasm where it utilizes 10 open reading frames (ORFs) to encode numerous non-structural proteins and 4 structural proteins [**9**]. As has been described for other coronaviruses, the four structural proteins are nucleocapsid (N), membrane (M), envelope (E), and spike (S) and are presumed responsible for maintaining the structural integrity of the enveloped SARS-CoV-2 virion [**9**]. The M glycoprotein of coronaviruses drives the assembly and formation of progeny viral particles from the endoplasmic reticulum-Golgi intermediary complex (ERGIC) and is the most abundant viral structural protein in the virion [**9**]. M oligomerizes to create a protein lattice across ERGIC membranes and interacts laterally with S and E, the other two viral membrane proteins, which are integrated into the structural matrix at budding sites [**3, 9**]. The role of E in assembly and budding is enigmatic, though it has been shown to be crucial for proper assembly of SARS-CoV-1 viral particles and serves as a viroporin altering ion transport [**3, 9, 10**].

The S protein gives coronavirus particles their pronounced crowned (‘corona’) structure [**9**]. While dispensable for viral particle assembly and formation in SARS-CoV-1, the incorporation of S is required for progeny viral particles to successfully infect a host cell [**2**]. The final structural protein, N, is responsible for coordinating the viral RNA genome to the structural matrix which it does through interactions with the cytosolic C-terminal endodomain of M in an RNA-independent manner [**3, 9**]. These interactions between the four structural proteins facilitate the proper assembly, genomic packaging and budding of progeny coronavirus particles [**9**]. After budding into the ERGIC lumen, progeny viral particles are released from the infected cell by exocytosis [**9**]. While these processes have yet to be fully examined specifically in the SARS-CoV-2 virus, they are likely to follow a similar scheme based upon homology.

Previously, transient co-expression of the four SARS-CoV-1 structural proteins in mammalian cell culture has been shown to produce self-assembling VLPs which can be collected, purified, and used to study the molecular biology of the virus [**2**]. Specifically, aspects of the viral lifecycle such as assembly [**3**], budding [**3, 11**], egress [**2, 11**], and entry [**12**] have been studied for SARS-CoV-1. While these VLPs were both morphologically and functionally similar to the authentic SARS-CoV-1 live virus, they do not contain the viral genome, are non-infectious, and thus can be used in a BSL-2 setting.

Until now, SARS-CoV-2 VLPs have only been used to identify M as the driver of viral particle formation [**13**] and for vaccine development [**14**]; they have yet to be functionalized to study SARS-CoV-2 entry or inhibitors. Instead, the few identified entry inhibitors of SARS-CoV-2 have been evaluated using classical coronavirus assays such as S-mediated cell-cell fusion [**15**] or pseudotyped VSV vectors [**16**]. VLPs offer a reliable and realistic model of S-mediated fusion and viral entry events in a BSL-2 setting. Herein, we discuss methods for production, purification, validation, and utilization of SARS-CoV-2 VLPs.

## Results

### SARS-CoV-2 VLP production is driven by M co-expression with N, E, or S

A previous study by Xu et. al showed that M was released into the media of HEK293T and Vero E6 cells at 48-hours post-transfection independent of other viral structural proteins [**13**]. We repeated this experiment, independently expressing M, N, E, or S in HEK293 cells and collecting VLPs at 72-hours post-transfection. Cell lysates and VLP fractions were analyzed with western blot analysis (**Figure 1A**). Additionally, we examined independently transfected cells with scanning electron microscopy (SEM) (**Figure 2, Figure S1**). When analyzed with western blot, M was not detectable in the cell lysate or VLP fraction when expressed independently. The insoluble cellular fraction was evaluated as well, and when expressed alone, M was also undetectable in this fraction (**Figure 1B**). However, when co-expressed with N or S, M was detectable in the cell lysate, insoluble, and VLP fraction (**Figure 1A, B, Figure S2**). This effect of N on M was confirmed with immunofluorescence (**Figure S3**). When analyzed with SEM, M independently expressed did appear to induce small changes to membrane structure but overall was qualitatively insignificant compared to mock transfections (**Figure 2**).

**Figure 1.**
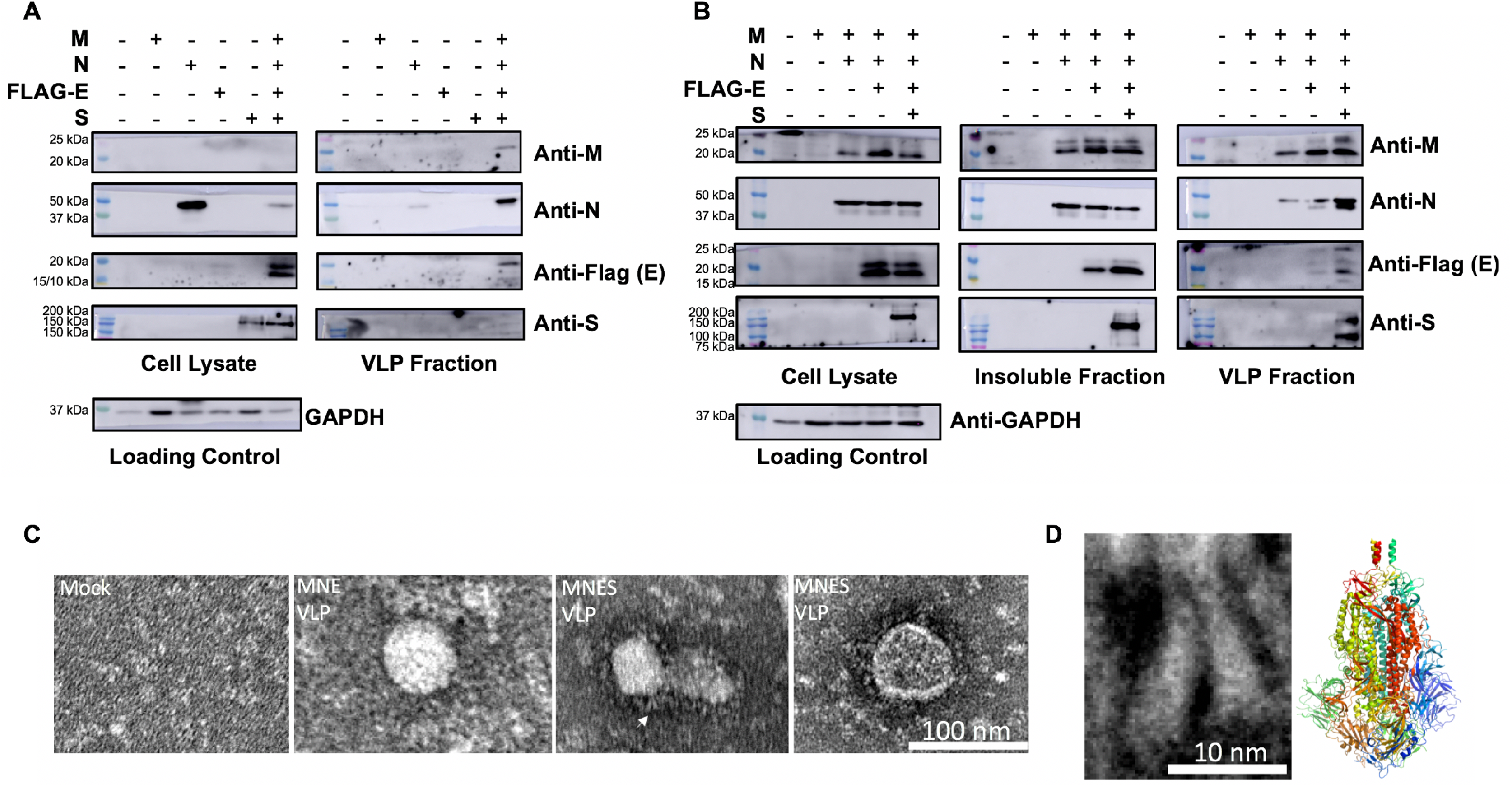
Production of SARS-CoV-2 virus-like particles (VLPs). **A**) Western blot analysis of the cell lysate and VLP fractions of both individual and full combination of the four structural proteins 72-hours post-transfection. Total protein content of cell lysates was used to normalize loading conditions and was quantified using the Pierce bicinchoninic acid assay. VLP loading was calculated as a constant ratio to normalized cell lysates. **B**) Western blot analysis of the cell lysate and VLP fractions of additive combinations of M, N, and S. Total protein content of the cell lysates was used to normalize loading conditions and was quantified using the Pierce bicinchoninic acid assay. VLP loading was calculated as a constant ratio to normalized cell lysates. **C**) Electron microscopy of SARS-CoV-2 VLPs. Purified M+N+E and M+N+E+S VLPs were added to glow discharged 400-mesh copper grids covered with carbon-coated collodion film. Grids were washed in one drop of water, stained in three drops of phosphotungstic acid (1.0% w/v), air dried, and imaged. White arrow demarcates zoomed image used for 1D. **D**) Two S molecules on the left insert are magnified from Figure 1C marked with white arrow. The right insert corresponds to cryo-EM structure of trimeric S protein (PBD: 6ZWV). Its longest dimension is ~170 angstroms which is comparable to negative staining S protein present around VLPs.

**Figure 2.**
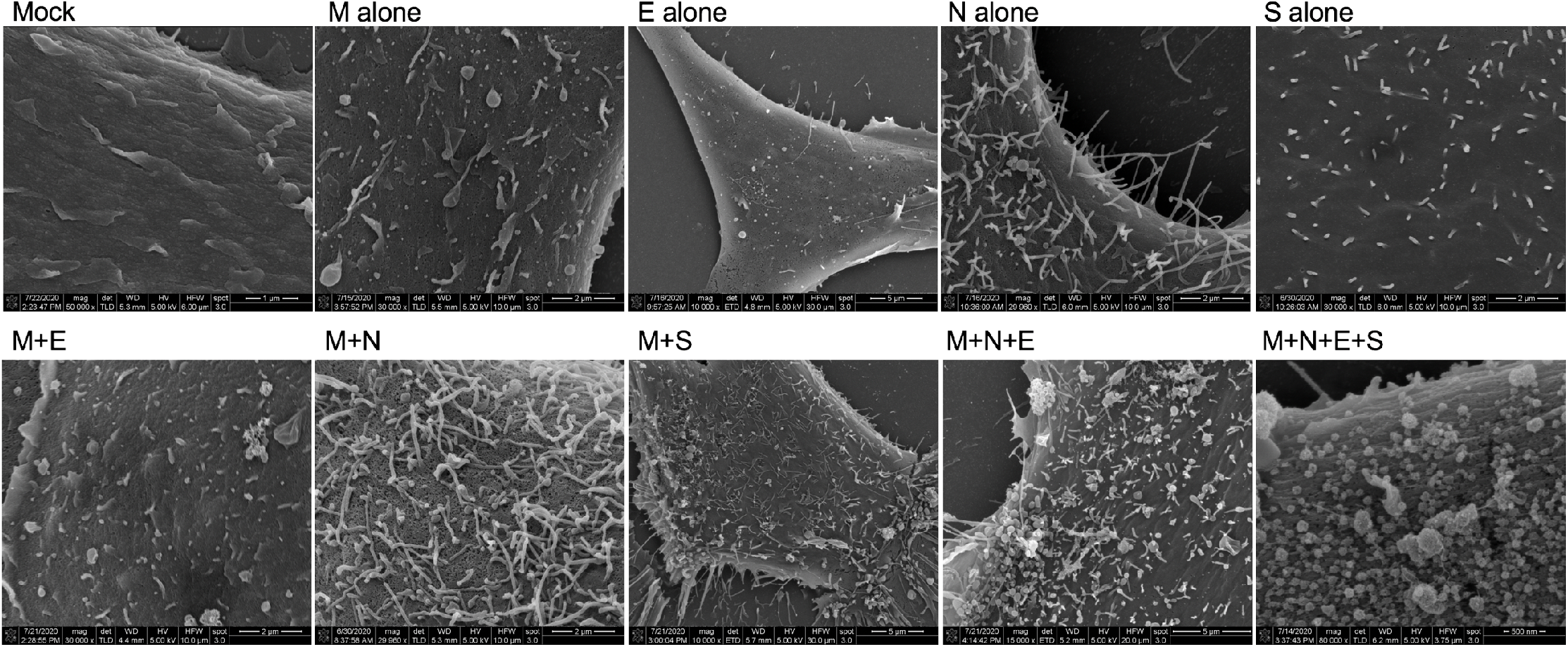
Scanning electron microscopy of viral structural protein transfected cells. HEK293 cells were seeded onto coverslips and transfected individually or in combination with M, N, E, and/or S. Cells were fixed with glutaraldehyde 72-hours post-transfection and kept at 4°C until fixed with osmium tetroxide. Samples were then gradually dehydrated with ethanol and completely dehydrated with a critical point dryer. Once dehydrated, samples were mounted onto aluminum pins with double-sided carbon tape, charged with silver paint, and sputter coated prior to imaging. Images range in magnification from 10,000x to 80,000x.

When independently expressed, N was readily detectable in the cell lysate and a small band was detectable in the VLP fraction (**Figure 1A**). Intriguingly, N was also found in the insoluble fraction (**Figure 1B**). When this VLP sample was analyzed with transmission electron microscopy (TEM), this fraction was not found to contain VLPs, but had proteinaceous aggregates that resemble high order N oligomers (**Figure S4**, white arrow). It is possible that what is detected in the western blot were either secreted N protein or detached N-packed filopodia which co-sedimented with VLPs, as SEM analysis revealed that expression of N alone induces filopodia formation. Further, filopodia formation in N-transfected cells was observed with confocal microscopy (**Figure S5**). These results are complimentary to a study which previously showed that filaments formed in SARS-CoV-2 infected cells are filled with N protein [**17**].

When independently expressed, E was not detectable in the cell lysate. Similar to M, E is seemingly stabilized by N as it becomes detectable in the cell lysate when co-expressed (**Figure 1B, Supporting Information 1**). Contradictory to Xu et al. and previous work studying other coronavirus VLPs [**13, 18, 19**], when E was independently expressed it was not detectable in the VLP fraction by western blot (**Figure 1A**). Further, TEM analysis of the VLP fraction revealed a proteinaceous background comparable to that of the mock transfected VLP collection. (**Figure S4**). SEM analysis of E-transfected cells revealed little to no change in the plasma membrane structure when compared to mock transfected cells (**Figure 2**).

When independently expressed, S was readily detectable in the cell lysate but not detectable in the VLP fraction. Interestingly, SEM analysis of S-transfected cells revealed stiff, rigid protrusions from the plasma membrane (**Figure 2**). S trafficking to the plasma membrane was confirmed by confocal microscopy using S-GFP (**Figure S6**).

Since none of the viral structural proteins alone could sufficiently support VLP formation, we cotransfected combinations of structural proteins with M, as M is thought to be the major driver of particle assembly in coronaviruses [**2, 9, 13**]. First, combinations of M+N (**Figure 1B**), M+E (**Figure S2**), and M+S (**Figure S2**) were co-transfected and VLPs collected 72-hours post-transfection. While M remained undetectable in the cell lysate and VLP fraction when expressed alone, when co-expressed with N or S, M was detectable in the cell lysate and VLP fraction. M+E did not release a detectable level of M. All three of these combinations were also examined with SEM (**Figure S1**). M+E did not produce major changes in plasma membrane structure; however, M+N and M+S did. In M+N transfected cells, long filamentous filipodia were observed while in M+S transfected cells, stiff, rigid filaments were observed.

We further explored triple combinations of M+N+E (**Figure 1B**) and M+E+S (**Figure S2**). M+N+E produced a detectable level of VLPs while M+E+S did not. The addition of E to the M+N system increased N incorporation and VLP production, indicating an important role of E in assembly and release. SEM analysis of M+N+E transfected cells revealed visible changes in membrane structure and VLPs at the cell surface. TEM analysis of the M+N+E VLP fraction revealed bald, spherical VLPs which were not observed in the mock sample (**Figure 1C**). When all four structural proteins were expressed and analyzed with TEM, spherical particles of approximately 100 nm in size were observed with a pronounced crown or ‘corona’ (**Figure 1C**). M+N+E+S transfected cells were also analyzed by SEM and revealed numerous VLPs at the cell surface, specifically at the base of filopodia. This is in agreement with SEM data examining release of SARS-CoV-1 viral particles [**20**] and recent SEM on SARS-CoV-2 infected cells [**17**]. When analyzed with western blot, the M+N+E+S condition released the most VLPs. These findings are again contradictory to those of Xu et al. where S seemed to limit M release; however, in their study all four structural proteins harbored a tag in contrast to our system.

### Transient production of VLPs can be used to model viral assembly and budding

To model VLP assembly and budding we utilized a recent electron microscopy technology, ascorbate peroxidase (APEX2) tagging, by exploiting S protein’s tolerance of a C-terminal tag to produce S-APEX2. APEX2 tagging works by catalyzing 3,3’-Diaminobenzidine (DAB) oxidation, which produces a dark brown precipitate visible with TEM [**21**].

For *in-situ* detection of VLPs, we co-expressed S-APEX2 with the other viral structural proteins M, N, and E. 30-hours post-transfection, cells were fixed and processed for cryo-APEX TEM. Cells exhibited localized staining of various compartments of the endomembrane system that extended from the perinuclear region nearly to the plasma membrane (**Figure 3A**, inset). At higher magnifications, images revealed gross perturbations of the endomembrane system with the appearance of highly stained localized vacuolar clusters enmeshed within tubular components (**Figure 3A**, white circles). The stained areas represent localization of the S-APEX2.

**Figure 3.**
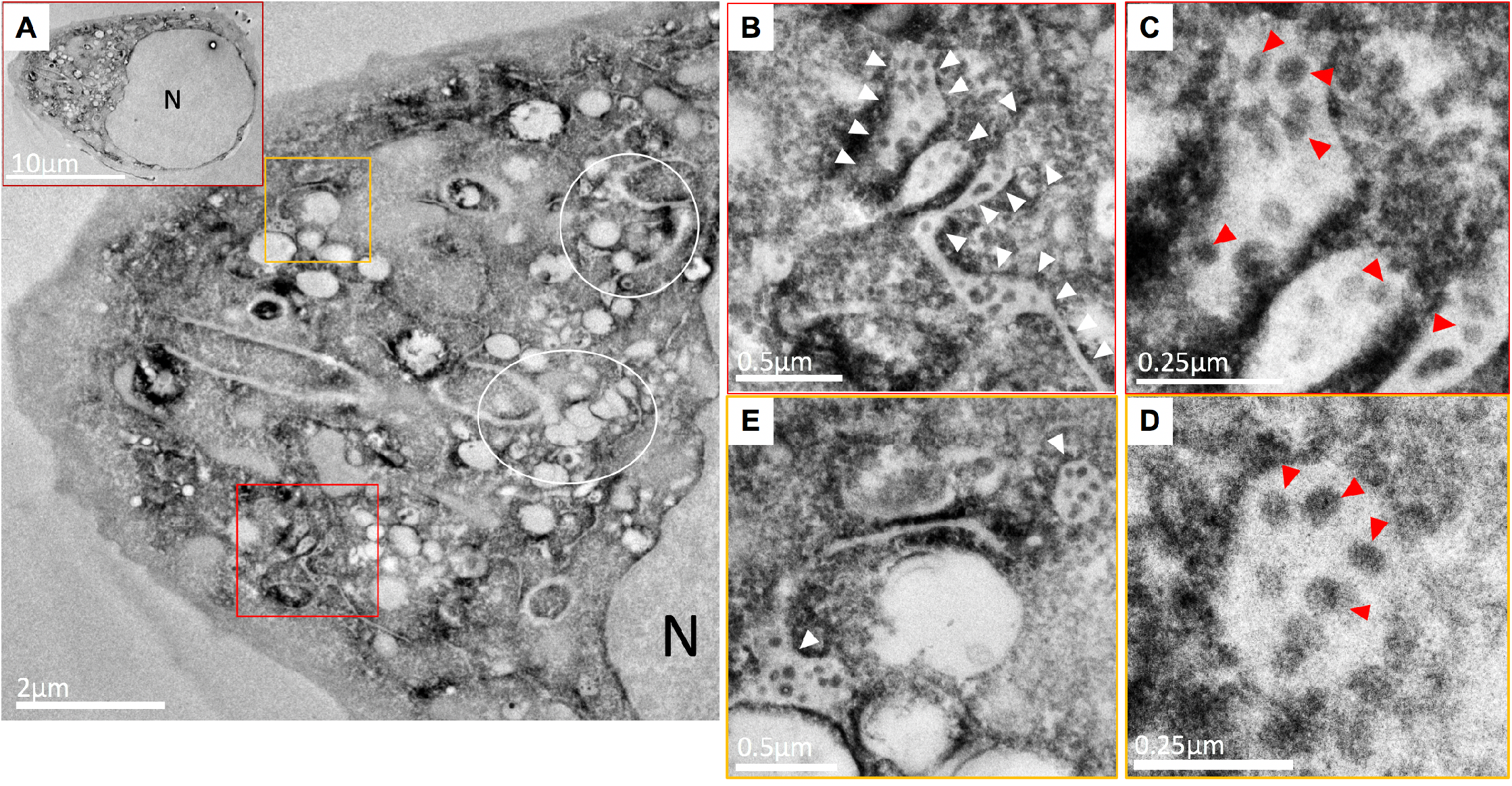
Detection and visualization of VLP assembly and budding via electron microscopy. M, N, E, and S-APEX2 were co-expressed in HEK293 cells. 30-hours post-transfection cells were fixed and processed for imaging. TEM images of ultra-thin sections of resin embedded cells subjected to APEX2-DAB assay exhibit intense staining of the perinuclear endomembrane system (A, inset). This region exhibited localized stained clusters of vacuolar structures enmeshed with tubular compartments (A, indicated with white rings). Some of these stained vesicular-tubular structures are filled with stained VLPs (A, within red and orange boxes). At higher magnification, they appear as swollen tubular structures resembling ERGIC compartments (B and E, white arrowheads) filled with stained spherical VLPs. The spherical structures within these compartments (C and D, red arrowheads) resemble VLPs in size range and morphology and carry the APEX2 specific stain.

VLPs were detected in some of the stained swollen vesicular-tubular structures that had the structural hallmark of the ERGIC (**Figure 3A**, marked by red and yellow squares). Demarcated areas revealed heavily stained tubular and vesicular compartments (**Figure 3B, E**, white arrowheads) that contained spherical VLPs evident due to the presence of APEX2-tagged spike protein (**Figure 3C,D**, red arrowheads). The VLPs in these compartments are well within the expected size range of SARS-CoV-2 virions. Spike protein alone cannot form VLPs, indicating the proper co-expression of the other three viral structural proteins.

### SARS-CoV-2 VLPs can be used to model viral entry

To produce fluorescently labeled VLPs, we co-expressed S-GFP with M, N, and E in HEK293 cells. S-GFP incorporation into VLPs (GFP-VLPs) collected 72-hours post-transfection was confirmed with western blot analysis (**Figure S7**). To test the entry competency of GFP-VLPs, the GFP-VLP entry assay was performed (**Figure 4A**, schematic). Target cells infected with GFP-VLPs had clear GFP signal present in punctate, intracellular structures while mock infected cells lacked detectable GFP signal (**Figure 4A**). As an additional control, media collected and clarified from cells expressing S-GFP alone was collected and used to infect target cells (**Figure S8**). Similar to mock infection conditions, media from S-GFP expressing cells failed to yield detectable GFP signal in infected target cells.

**Figure 4.**
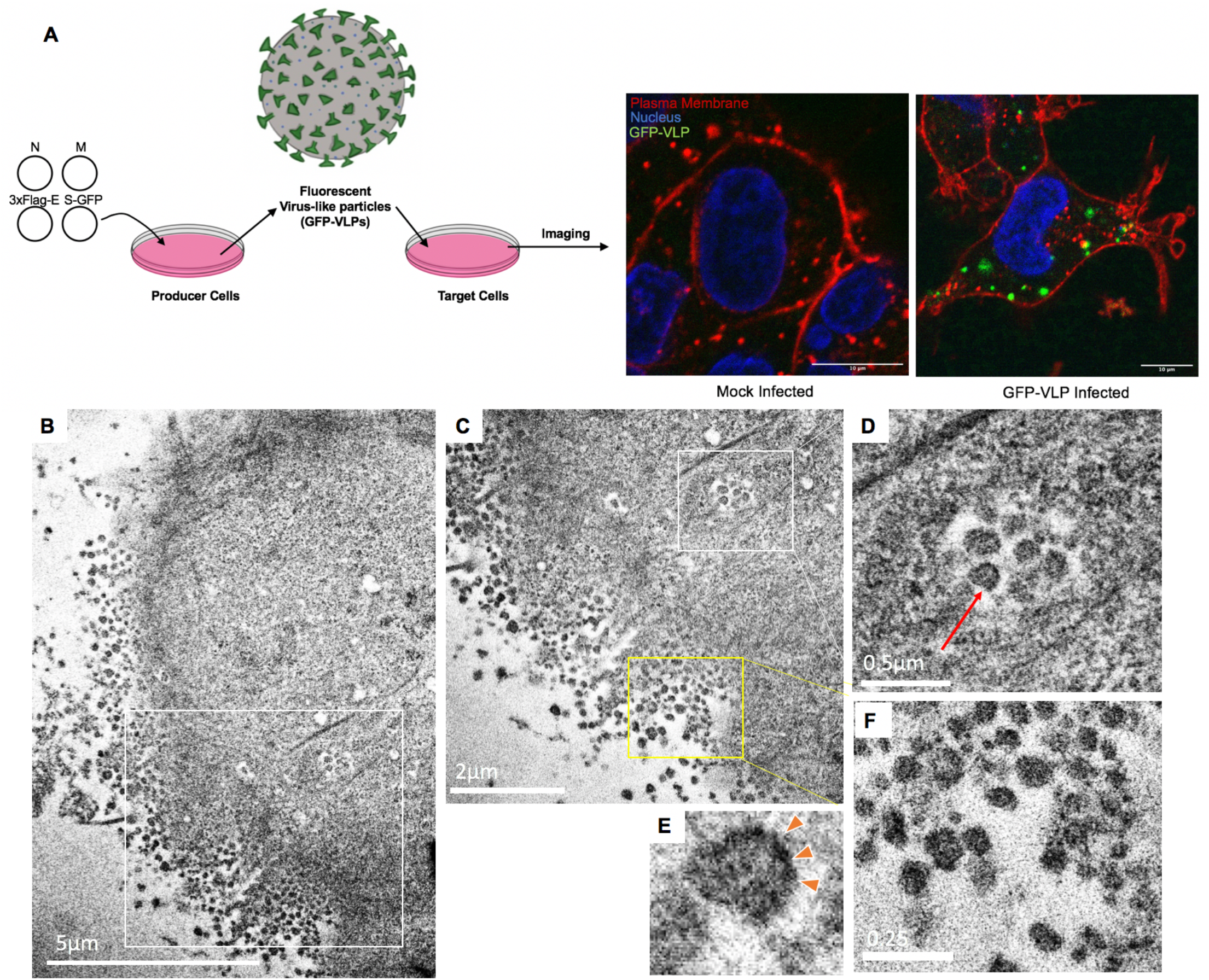
Detection of VLP entry of target cells with confocal and electron microscopy. A) Schematic of the GFP-VLP entry assay. GFP-VLPs were produced in HEK293 cells and used to infect target cells. After spinoculation and 2-hour incubation, cells were fixed, stained with plasma membrane (WGA-Alexa647) and nuclear (Hoechst 3342) stains, and imaged on the confocal microscope. B) APEX-VLPs were produced in HEK293 cells and used to infect target cells. After spinoculation and 2-hour incubation, cells were fixed, processed, and imaged with TEM. APEX reaction was performed on coverslips and blocks sectioned *en face* to be able to image stained VLPs at the cell periphery (B, and magnified image of the area within white box in A, shown in C and F). At this stage, endosomes filled with stained VLPs are observed entering the cell (indicated with a white box in C and at higher magnification in D). E) At higher magnification, these stained structures are seen to have a darker stained periphery as expected of VLPs with stained S protein (D, red arrow and magnified image of the same in E). F) Magnified image of the stained VLPs at the periphery of the cell show a size distribution that falls within the expected range reported for SARS-CoV-1 and SARS-CoV-2 virus.

To model viral entry using TEM, we utilized S-APEX2 incorporated VLPs. Using the same methodology from the GFP-VLP entry assay, an APEX-VLP entry assay was performed. Infected target cells were fixed, processed, and imaged with TEM as previously described in assembly and budding (**Figure 4B-F**). Dark staining represents the APEX2 signal and thus the localization of the APEX-VLPs. They were clearly detected clustered in large internalized vesicles at the periphery of the cell.

### VLP entry correlates with authentic live virus entry

Coronaviruses in general are understood to utilize the endocytic pathway to gain entry into target cells [**9**]. Recently, SARS-CoV-2 viral particles were shown to colocalize with endocytic markers early endosome antigen 1 (EEA1) and lysosomal associated membrane protein 1 (LAMP1) after a 90-minute incubation with target cells [**22**]. Viral particles were more frequently colocalized with LAMP1+ vesicles than EEA1+ vesicles, indicating that at 90-minutes post-infection, most particles are already trafficked deep into the endocytic pathway.

To evaluate the entry mechanism of our SARS-CoV-2 GFP-VLPs and compare it to that of the live virus, we performed the GFP-VLP entry assay on target cells pre-transfected with endocytic pathway markers mCherry-Rab5 (early endosomes) and mCherry-LAMP1 (lysosomes) (**Figure 5A**). Images were subsequently analyzed for green/red colocalization using the JACoP plug-in for ImageJ to calculate Pearson’s coefficient (**Figure 5B**). Pearson’s coefficient can be used to measure colocalization between two channels with values ranging from 1 for perfectly correlated fluorescence intensities to −1 for perfectly, but inversely, related fluorescence intensities. Near 0 are values which reflect two channels with intensities that are uncorrelated to each other [**23**].

**Figure 5.**
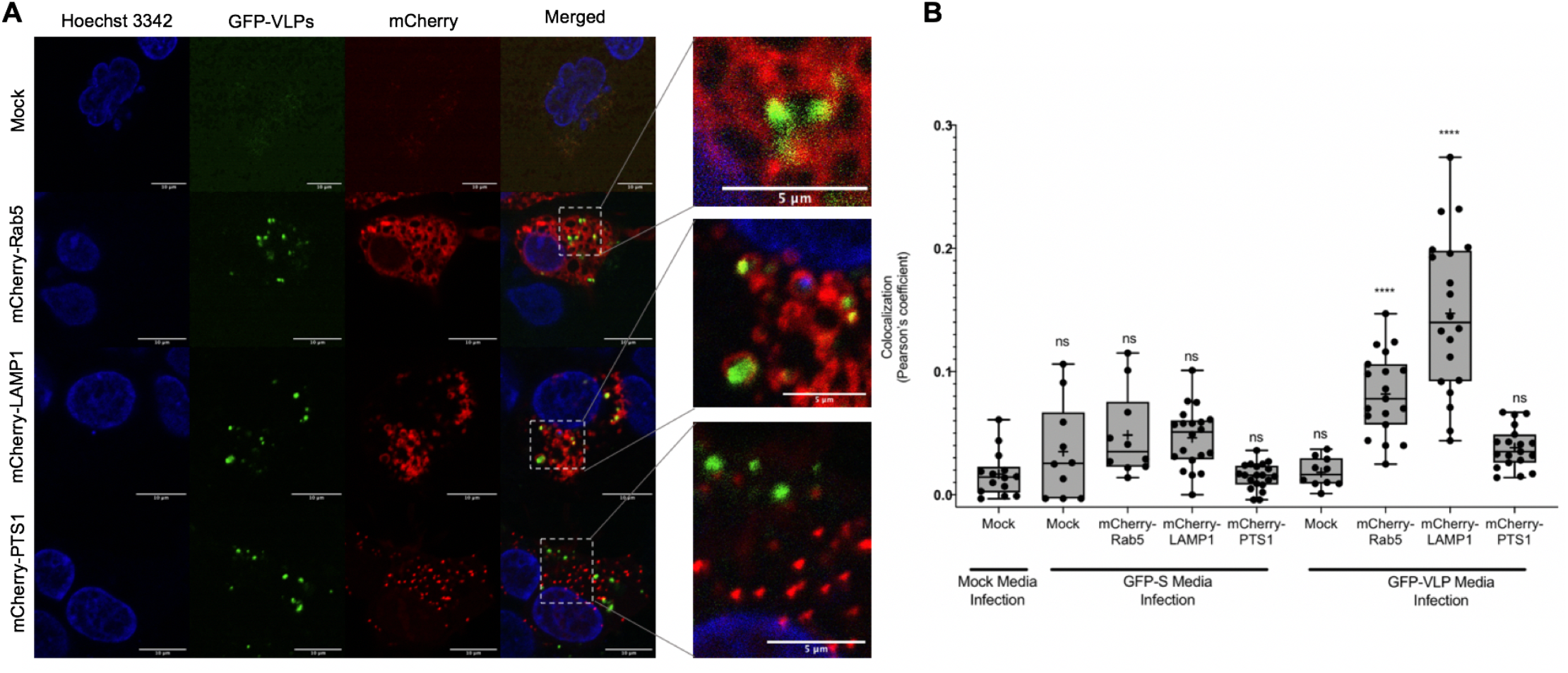
Colocalization of GFP-VLPs with endocytic markers. **A**) GFP-VLPs were produced and used to infect target cells co-expressing hACE2 and an mCherry-tagged marker for early endosomes (Rab5), lysosomes (LAMP1) or peroxisomes (PTS1). After infection, target cells were stained with Hoechst 3342 nuclear stain, fixed with 4% PFA, and then imaged with confocal microscopy. **B**) Images were analyzed for green/red colocalization using the JACoP plug-in for ImageJ to calculate Pearson’s coefficient and graphed with Prism. Statistics were calculated using an ordinary one-way ANOVA with multiple comparisons to mock infection of mock transfected target cells. GFP-VLP infection of mCherry-Rab5 and mCherry-LAMP1 was found to have significant colocalization (****) between the GFP and mCherry channels.

As controls, media from mock and S-GFP expressing producer cells were used to infect mock or endocytic marker-expressing target cells which were imaged and analyzed for green red colocalization. Colocalization analysis shows no significant colocalization between GFP and mCherry signals under any of these negative control conditions. When GFP-VLPs were used to infect mCherry-LAMP1 expressing target cells, GFP and mCherry signals colocalized with an average Pearson’s coefficient of 0.137, which was statistically significant when compared to GFP-VLP infection of mock transfected cells (0.021). When used to infect mCherry-Rab5 transfected target cells, GFP-VLPs and early endosomes had an average colocalization of 0.079, which was also statistically significant when compared to mock. When used to infect mCherry-PTS1 transfected target cells, GFP-VLPs and peroxisomes had an average colocalization of 0.036, which was not statistically significant when compared to the mock. Thus, SARS-CoV-2 VLPs localized with Rab5 positive and LAMP1 positive puncta as previously reported for authentic SARS-CoV-2. These VLP systems represent a novel approach for examining SARS-CoV-2 entry mechanisms in a BSL-2 setting.

## Discussion

As SARS-CoV-2 continues to spread, it is imperative that we continue to grow our fundamental understanding of its molecular virology. In this work, we examined the ability of viral structural proteins to produce VLPs and found that M alone was not sufficient to support VLP formation, but co-expression with N or S was the minimal requirement for VLP formation. E protein was found to have an additive effect in both N incorporation into VLPs and VLP production, highlighting the important role E must play in viral assembly and release. Additionally, this highlights the need for further examination of the role of E during infection. Finally, addition of S increased N and E incorporation further and increased VLP production, suggesting that M+N+E+S is most efficient at VLP production.

These findings are partially in contrast to the findings of Xu et. al [**13**]; however, the controversy over the minimal efficient system for SARS-CoV-2 VLP production is paralleled by the controversy over SARS-CoV-1 VLP production. One study of SARS-CoV-1 VLPs suggests the minimal requirement for efficient VLP production was M+E [**24**], while another study showed the minimal system was M+N [**2**]. Additionally, Siu et. al showed that the most efficient system for SARS-CoV-1 VLP production was M+N+E. As for MERS and other coronaviruses such as mouse hepatitis virus, bovine coronavirus, infectious bronchitis virus, and transmittable gastroenteritis virus, M+E was found to be the minimal system for efficient VLP production [**2, 25**].

While N was shown to be important in increasing VLP formation, in this work we also show that N drives the formation of filamentous filopodia in transfected cells. These findings compliment authentic live virus data which recently detected the formation of filopodia in SARS-CoV-2 infected cells [**17**]. It is hypothesized that these filopodia help progeny virus particles travel to and infect adjacent cells, which is supported by our SEM imaging that revealed large numbers of viral particles released at the base of filopodia. Taken together with data showing N in the insoluble fraction of HEK293 cells, this suggests that N may have lipid-binding properties; something that we plan to address in future studies.

In this work we also present for the first time a realistic model of SARS-CoV-2 viral entry available in a BSL-2 setting: SARS-CoV-2 GFP- and APEX2-VLPs. In accordance with live virus data, GFP-VLPs colocalize with the early endosome marker, Rab5, and the late endosome marker, LAMP1. In future work, we plan to miniaturize our GFP-VLP entry assay and use it to screen for viral uptake and entry inhibitors. Not only is confocal microscopy available for evaluation of GFP-VLP entry events, we utilized APEX tagging technology to make evaluation of SARS-CoV-2 entry accessible to electron microscopy.

Traditionally, TEM has been used to demonstrate VLP-like structures in large vacuoles in cells transfected with plasmids encoding structural proteins; however, many times such identification was based solely on morphology [**26**]. By utilizing APEX tagging, we have shown for the first time localization of S protein during VLP assembly and budding as well as the formation and export of APEX-VLPs from the presumed ERGIC lumen. In total, this research provides ample resources for other BSL-2 laboratories interested in joining the growing field to try and understand SARS-CoV-2 assembly, budding, and entry dynamics, biochemical and biophysical questions on the four structural proteins, and drug screening of viral assembly, budding, and/or entry inhibitors.

## Experimental Procedures (Materials & Methods)

### Plasmid constructs

The pcDNA3-Membrane, pcDNA3-HA-Membrane, pcDNA3-Nucleoprotein, and pCMV 3xFlag-Envelope plasmids were a kind gift from Erica Sapphire (The La Jolla Institute of Immunology, La Jolla, CA). The pCAGGS-Spike plasmid was from BEI Resources (NR-52310). The pcDNA3.1 Spike-GFP plasmid (Genescript, MC_0101089) was a generous gift to us by Raluca Olstafe (Purdue University). The pcDNA3.1-Spike-APEX2 plasmid was synthesized by Gene Universal (Newark, DE, USA). mCherry-Lysosomes-20 (i.e. LAMP1, Addgene #55073), mCherry-Rab5a-7 (Addgene #55126), and mCherry-Peroxisomes (Addgene #54520 were gifts from Michael Davidson. pcDNA3.1-hACE2 was a gift from Hyeryun Choe (Addgene plasmid # 1786) of Scripps Research, Florida [**27**].

### Cells and culture conditions

Human embryonic kidney (HEK293) cells (from American Type Cell Collection, Manassas, VA, USA) were maintained in Dulbecco’s modified Eagle’s medium (DMEM) supplemented with 10% fetal bovine serum, 1% penicillin-streptomycin, and 1% MEM Non-essential amino acids maintained at 37°C and 5% CO_2_ conditions.

### Transient Transfections

Transfections were performed using 2.5 M CaCl_2_ and 2X HBS [10 mM D-Glucose, 40 mM HEPES, 10 mM KCl, 270 mM NaCl, 1.5 mM Na_2_HPO_4_, pH=7.06] as published by Abcam [**28**]. Briefly, DNA was mixed in ddH_2_O, CaCl_2_ was added, and the tube was mixed lightly. 2X HBS was then added to the tube for 1X final concentration, dropwise. Subsequently, the solution was mixed well and incubated at room temperature for 20 minutes. Transfection mixtures were added dropwise on to cells in DMEM + 10% FBS and incubated for the specified time.

### Production of SARS-CoV-2 VLPs

HEK293 cells 70% confluent in 100mm dishes were transfected with 6 μg of each SARS-CoV-2 pcDNA3-M, pcDNA3-N, pCVM-3xFlag-E, and/or pCAGGS-S. Transfection mixtures were prepared in ddH20 using 100 ng/μL stock DNA combined with transfection reagents 2.5M CaCl_2_ and 2X Hepes buffered saline, incubated for 20 minutes at room temperature, then added dropwise to cells in DMEM + 10% FBS + 1% PS + 1% MEM Non-essential amino acids. Cells used for the production of VLPs are referred to as “producer cells”.

### Purification of SARS-CoV-2 VLPs

Similar to methods described for the purification of SARS-CoV VLPs [**2**], SARS-CoV-2 VLPs were purified from the media of producer cells 72-hours post-transfection. Media was removed from cells and clarified with light centrifugation at 1,000 x g for 10 minutes at room temperature. Clarified VLP-containing media was loaded on top of a 20% sucrose cushion using a glass pipette and then ultracentrifuged for 3 hours at 4°C and 100,000 x g in a Beckman Type 70 Ti rotor. VLP-containing pellets were carefully resuspended in TNE buffer (50mM Tris-HCl, 100 mM NaCl, 0.5 mM EDTA, pH=7.4) containing 5% sucrose. Cell lysates of producer cells were prepared using 500μL RIPA buffer containing 1X Halts protease inhibitor and 0.5% N-Lauroylsarcosine for 1 hour on ice, vortexing every 15 minutes, followed by a 10-second sonication and finally ultra-centrifugation at 25,000xG for 20 minutes at 4°C; the soluble fraction was collected and the pellet was resuspended in 500μL RIPA buffer containing 1X Halts protease inhibitor and 0.5% N-Lauroylsarcosine by sonicating 10 seconds.

### Western blot analysis of VLPs and cell lysates

Total protein content of producer cell lysates was used to normalize loading conditions and was quantified using the Pierce bicinchoninic acid assay (BCA). VLP loading was calculated as a constant ratio to normalized cell lysates. Samples were prepared in 1X E running buffer (5M Urea, 1M DTT, 100mM NaCO_2_, 25mM Tris, 0.5% CHAPS, pH=11) with 1X reducing Laemmli SDS Sample buffer (Alfa Aesar), resolved in a 10% SDS-PAGE polyacrylamide gel, and subsequently transferred on to a supported 0.45 μm nitrocellulose membrane (BioRad) which was used for immunoblotting. Membranes were cut and probed for M with rabbit anti-SARS-1 M (Rockland), N with rabbit anti-SARS-2 N (Genetex), 3xFlag-E with mouse anti-Flag IgG HRP (Abcam), and S with mouse anti-SARS1/2-Δ10S [1A9] (Genetex); goat anti-rabbit IgG HRP (Abcam) and sheep anti-mouse IgG HRP (Abcam) secondary antibodies were used as appropriate. GAPDH was used as a cell lysate loading control by probing membranes with mouse anti-GAPDH monoclonal IgG (Thermofisher Scientific) followed by sheep anti-mouse IgG HRP. All antibodies were diluted in 5% milk and membranes were washed with 1X TBST. Chemiluminescent signal was visualized using Clarity™ Western ECL Substrate (BioRad) or Clarity Max Western ECL Substrate (BioRad) and imaged using an Amersham Imager 600 (GE Healthcare Life Sciences).

### Immunofluorescence

HEK293 cells of 70% confluency grown in the 8-well glass bottom plates were transfected with pcDNA3-N using either lipofectamine LTX or 2000 reagent (Thermofisher scientific) according to the manufacturer’s instructions. At 24-hours post-transfection, the cells were fixed by 4% paraformaldehyde (PFA), permeabilized by 0.1-0.2% Triton X-100 in PBS, blocked by 2.5-3% FBS and 1% BSA in PBS and then incubated with the appropriate primary antibody overnight (Rabbit anti-SARS-CoV-2-N antibody, GeneTex #GTX135357, 1:500 or Rabbit anti-SARS-1-M, Rockland, 1:1000; mouse anti-GORAPS2 (Golgi marker), Sigma Aldrich, 1:500). Following overnight incubation, cells were washed and then incubated with the appropriate secondary antibody (anti-rabbit IgG-Atto 594, Millipore Sigma #77671-1ML-F, 1:1000; anti-mouse IgG-Atto488, Millipore, 1;10000) for 45 min – 1 hour at room temperature. Following secondary incubation, cells were washed, stained with Hoechst 3342 nuclear stain, and then imaged with confocal microscopy. Images were analyzed using ImageJ. For images of N protein alone, some cell body areas were saturated to achieve filopodia visibility.

### Negative staining transmission electron microscopy

To prepare the grids for negative stain EM, 4 μl of purified VLPs were added to a glow discharged 400-mesh copper grid covered with carbon-coated collodion film (EMS, Hatfield, PA). Grids were washed in one drop of water, stained in three drops of Phosphotungstic acid (1.0% w/v) (EMS, Hatfield, PA) and air dried. Samples were visualized on a Tecnai G2 T20 electron microscope (FEI, Hillsboro, OR) at an acceleration voltage of 200 kV. Images were taken at a magnification of 43,000× at a defocus value of −1.4 μm and recorded on a Gatan US1000 2Kx2k CCD camera (Gatan, Pleasanton, CA). Images were converted to mixed raster content format, resulting in final images with a pixel size of 4.23 Å/pixel at the specimen level.

### Scanning electron microscopy

Silica coverslips were placed on the bottom of 12-well plates before being seeded with HEK-293 cells to 30% confluency for transfection 24-hours after seeding. Calcium chloride transfection was conducted with structural protein cDNA vectors and cells were incubated in Dulbecco’s Modified Eagle Media (DMEM) plus 10% Fetal Bovine Serum (FBS) for 72 hours. At 72 hours post transfection, cells were fixed with primary fixative, 2.4% glutaraldehyde, 0.1 M cacodylate fixative buffer and sealed in parafilm at 4°C. Samples were then processed by the Purdue Electron Microscopy Facility, washing coverslips with 0.1 M cacodylate buffer before the addition of secondary fixative, 4% osmium tetroxide, 0.1 M cacodylate buffer and incubated for 30 minutes. Samples were then dehydrated gradually with increasing percentage of ethanol and then dried in the critical point dryer (Tousimis AutoSAMDRI-931, CPD) available at the facility. After drying, coverslips were mounted onto aluminum pin stub mounts with double-sided conductive tape, conductive liquid silver paint, and sputter coated for 60 seconds. Upon completion of the samples were visualized and imaged on the FEI Nova NanoSEM at the Purdue Life Science Electron Microscopy Facility.

### GFP-VLP Entry Assay

Following the protocol outlined in *Production of SARS-CoV-2 VLPs*, SARS-CoV-2 pcDNA3-M, pcDNA3-N, pCVM-3xFlag-E, and pcDNA3.1-S-GFP were co-expressed in HEK293 cells. 72-hours post-transfection, media was collected from producer cells and clarified with light centrifugation at 1,000xG for 10 minutes at room temperature. 5 mL of clarified media was added per well to 70% confluent target HEK293 cells plated in a black, glass-bottom 6-well plate (Cellvis, Mountain View, CA, USA). Target cells were then spinoculated using an M-20 microplate swinging bucket rotor at 2,000 rpm, 4°C, for 1 hour. After spinoculation, target cells were placed at 37°C for 2 hours. For imaging-based experiments, following this incubation infected target cells were stained with Hoechst 3342 nuclear stain and wheat germ agglutinin (WGA)-Alexa 647 plasma membrane stain, fixed with 4% paraformaldehyde for 7 minutes at room temperature, and then imaged with confocal microscopy.

### APEX-VLP Entry Assay

Following the protocol outlined in *Production of SARS-CoV-2 VLPs*, SARS-CoV-2 pcDNA3-M, pcDNA3-N, pCVM-3xFlag-E, and pcDNA3.1-S-APEX2 were co-expressed in HEK293 cells. 72-hours post-transfection, media was collected from producer cells and clarified with light centrifugation at 1,000 x g for 10 minutes at room temperature. 5 mL of clarified media was added per well to 70% confluent target HEK293 cells transiently expressing hACE2 plated on 22mm glass coverslips in a 6-well plate. Target cells were then spinoculated using an M-20 microplate swinging bucket rotor at 2,000 rpm, 4°C, for 1 hour. After spinoculation, target cells were placed at 37°C for 2 hours. Post-incubation, cells were fixed with 2.5% glutaraldehyde in 0.1% sodium cacodylate buffer (pH 7.4) were kept on ice for 30 minutes. Cells were kept between 0 and 4°C for all subsequent steps until resin infiltration. Cells were washed 5 times, 3 minutes each, with chilled cacodylate buffer and incubated in 1 mL of freshly prepared 0.5 mg/mL 3,3’-diaminobenzamide (DAB) (Sigma-Aldrich) for 2 minutes. DAB combined with 10mM of hydrogen peroxide was then added to the cell and incubated for 15 minutes or until a dark brown color develops. DAB was removed and the cells were washed 3 times, 5 minutes each, followed by staining with 1% Osmium tetroxide for 10 minutes on ice. Cells were washed 2 times, 5 minutes each, in chilled cacodylate buffer and twice with water. Cells were then dehydrated in a graded ethanol series (50%, 70%, 90%, 95%, 100%, 100%, 100%), for 10 minutes each and then infiltrated with an increasing concentration of Durcupan ACM resin (Sigma-Aldrich) in ethanol (30%, 60%, 90% 2 hours each and then overnight in 100% and 2 hours twice in 100%+ component C) with gentle rocking. The coverslips with cells on them were then picked up with tweezers and planted face down on BEEM^®^ capsules (Electron Microscopy Sciences) pre-filled with 100%+C and baked in the oven at 60°C for 36 hours. Coverslips were separated from the BEEM^®^ capsules by dipping them in liquid nitrogen. Blocks were then extracted from the BEEM^®^ capsules, loaded onto the ultramicrotome. 90nm sections were obtained *en face* from the single layer of cells using a diamond knife (DiATOME) and imaged on a T12 (FEI) transmission electron microscope.

### TEM-based subcellular localization of VLP assembly using CryoAPEX

Subcellular localization of tagged VLPs were visualized using the CryoAPEX method as described in Sengupta et. al [**29**]. Following the protocol outlined in *Production of SARS-CoV-2 VLPs*, SARS-CoV-2 pcDNA3-M, pcDNA3-N, pCVM-3xFlag-E, and pcDNA3.1-S-APEX2 were co-expressed in HEK293 cells. At 48-hours post-transfection, cells were dislodged from the plates using trypsin, pelleted at 500 x g and fixed with 2.5% glutaraldehyde. The pellet was then washed 3 times for 5 minutes each with chilled 0.1 M sodium cacodylate buffer. Cells were subsequently incubated in a combination with 10mM of hydrogen peroxide and 1mg/mL of 3,3’-diaminobenzamide (DAB) in sodium cacodylate buffer, pH 7.4 for 30 minutes. Cells were then pelleted and washed 3 times using chilled sodium cacodylate buffer and then subjected to high pressure freezing (Leica, EM PACT 2) followed by freeze substitution (Leica) in the presence of 0.5% osmium tetroxide and 5% water in acetone. Cell pellets were then embedded in resin blocks and baked at 60°C for 36 hours. Thin (90nm) serial-sections were collected on formvar coated grids obtained on a microtome (Leica) using a diamond knife (DiATOME) and imaged on a T12 (FEI) TEM operating at 80kV.

### GFP-VLP Colocalization with Endocytic Markers

Following the protocol outlined in *GFP-VLP Entry Assay*, SARS-CoV-2 GFP-VLPs were prepared in HEK293 cells. 72-hours post-transfection, media was collected from producer cells and clarified with light centrifugation at 1,000 x g for 10 minutes at room temperature. 5 mL of clarified media was added per well to 70% confluent target HEK293 cells plated in black, glass-bottom 6-well plates (Cellvis, Mountain View, CA, USA), which had been transfected approximately 16-hours previously with 1μg hACE2 and 1μg mCherry-LAMP1, 1μg hACE2 and 1μg mCherry-Rab5, 1μg hACE2 and 1μg mCherry-PTS1, or mock. Target cells were then spinoculated using a M-20 microplate swinging bucket rotor at 2,000 rpm, 4°C, for 1 hour. After spinoculation, target cells were placed at 37°C for 2 hours. Following this incubation, cells were stained with Hoechst 3342 nuclear stain and wheat germ agglutinin (WGA)-Alexa 647 plasma membrane stain, fixed with 4% paraformaldehyde for 7 minutes at room temperature, and then imaged with confocal microscopy.

### Fluorescence microscopy

All confocal imaging was performed using the Purdue College of Pharmacy Live Cell Imaging Facility Nikon Eclipse Ti A1 instrument using NIS-elements AR software to capture 1024×1024 pixel resolution images at ¼ frame/second on 60X oil objectives detecting the fluorophores with channels in series.

## Acknowledgements

In addition to all of the authors of this manuscript, we would also like to acknowledge and thank Dr. Nathan Dissinger for his help in maintaining cells and preparing plasmid DNA. We would like to acknowledge the Purdue Life Science Electron Microscopy and Pharmacy Life Cell Imaging Facility for letting us use their facilities for support of imaging performed in this work.

